# Mechanisms involved in the active secretion of CTX-M-15 β-lactamase by pathogenic *E. coli* ST131

**DOI:** 10.1101/2021.03.31.437630

**Authors:** Severine Rangama, Ian D. E. A Lidbury, Jennifer M. Holden, Chiara Borsetto, Andrew R. J. Murphy, Peter M. Hawkey, Elizabeth M. H. Wellington

## Abstract

Infections caused by antimicrobial resistant bacterial pathogens are fast becoming an important global health issue. Strains of *Escherichia coli* are common causal agents of urinary tract infection and can carry multiple resistance genes. This includes the gene *bla*_CTX-M-15_ that encodes for an extended spectrum beta-lactamase (ESBL). While studying antimicrobial resistance (AMR) in the environment we isolated several strains of *E. coli* ST131 downstream of a WWTP in a local river. These isolates were surviving in the river sediment and characterisation proved that a multi-resistant phenotype was evident. Here, we show that *E. coli* strain 48 (river isolate ST131), provided a protective effect against a third-generation cephalosporin (cefotaxime) for a susceptible *E. coli* strain 33 (river isolate ST3576) through secretion of a functional ESBL into the growth medium. Furthermore, extracellular ESBL activity was stable for at least 24 h after secretion. Proteomic and molecular genetic analyses identified CTX-M-15 as the major secreted ESBL responsible for the observed protective effect. In contrast to previous studies, OMVs were not the sole route for CTX-M-15 secretion. Indeed, mutation of the Type I secretion system led to a significant reduction in the growth of the ESBL-producing strain as well as a significantly reduced ability to confer protective effect. We speculate that CTX-M-15 secretion, mediated through active secretion using molecular machinery provides a public goods service by facilitating the survival of otherwise susceptible bacteria in the presence of cefotaxime.

**Abstract importance:** Infections caused by antimicrobial resistant bacterial pathogens have become an important global health issue. Wastewater treatment plants (WWTPs) have been identified as hotspots for the dissemination of antimicrobial resistant genes/bacteria into the environment. In this study, we investigated resistance enzyme secretion by a multi-drug resistant human pathogenic *E. coli*, isolated from a UK river, downstream of a WWTP. We present evidence that the resistant strain actively secreted an important resistance enzyme into the surrounding medium which degraded the antibiotic cefotaxime. This research provided evidence for the mechanism for secretion of this enzyme which could indicate a new target to tackle antibiotic resistance pathogens.

## Introduction

Pathogenic strains of *Escherichia coli* producing CTX-M β-lactamases have recently emerged worldwide and now present the most common type of extended-spectrum β-lactamase (ESBL) enzymes in *Enterobacteriaceae* (Canton and Coque, 2006; Coque *et al.*, 2008; Rossolini *et al.*, 2008; Bevan *et al.*, 2017; Pitout and DeVinney, 2017). Limited treatment is available for patients infected with these *E. coli* strains which presents severe challenges to healthcare (Coque *et al.*, 2008; Coque *et al.*, 2008a; Hawkey and Jones, 2009; Bush and Jacoby, 2010; Peirano and Pitout, 2010; Bevan *et al.*, 2017). The global emergence of CTX-M producing *E. coli* is driven by the rapid dissemination of the gene *bla*_CTX-M_ located on highly mobilizable elements such as plasmids and transposons (Canton and Coque, 2006; Woodford *et al.*, 2009). Over 80 variants of CTX-M have been identified which cluster into five groups, CTX-M-1, -2, -8, -9 and -25 groups (Poirel *et al.*, 2005). The *bla*_CTX-M-15_ belongs to Group 1 and is the predominant variant in the human population globally, including the UK (Canton and Coque, 2006; Hawkey and Jones, 2009).

Clinical studies have suggested that secretion of hydrolytic enzymes such as β-lactamases irreversibly inactivate antibiotics outside the cell thus protect both the producer and otherwise susceptible bacteria in close proximity (Brook, 2004). One proposed mechanism for the secretion of ESBLs is the formation and likely stochastic release of outer-membrane vesicles (OMVs), which are common in Gram-negative bacteria (Mashburn-Warren and Whiteley, 2006). This secretory process eliminates the need for bacterial contact, or complex molecular architectures at the cell wall-periplasm interface typically required for long distance dissemination of extracellular proteins (Kulp and Kuehn, 2010). The packaging of β-lactamases into OMVs has been demonstrated in *Pseudomonas aeruginosa* by microscopy and enzymatic studies (Ciofu *et al.*, 2000). In addition, the release of OMVs containing various antibiotic-related proteins from a drug-resistant resistant *E. coli* facilitated the survival of various susceptible bacteria in presence of β-lactam antibiotics (Kim *et al.*, 2018). However, in this study the relative contribution of ESBLs compared to other antibiotic-related proteins, such as bacterial transporter systems, was not conclusively determined. Thus, the mechanism of CTX-M variants, such as CTX-M-15, in providing a protective effect remains uncertain. In Gram-negative bacteria, secretion of extracellular enzymes is achieved through either a one- or two-step process. In the two-step process, initial translocation across the cytoplasmic membrane to the periplasm is achieved through two main pathways: the twin-arginine (Tat) or the general secretory (Sec) pathway. Whilst the Tat pathway translocates a small number of folded proteins across the cytoplasmic membrane, the majority of proteins are translocated in an un-folded state using the Sec pathway (Robinson *et al.*, 2011). The second translocation event across the outer membrane is coordinated by various specialized export systems classified as Type II and Type V secretion systems (T2SS and T5SS, respectively). The one-step process, performed by Type 1 (T1SS), Type III (T3SS), Type IV (T4SS), and Type VI (T6SS) secretion systems, translocate proteins directly from the cytoplasm to the extracellular milieu, by passing the periplasmic space (Green and Mecsas, 2016; Tsirigotaki *et al.*, 2017). The architecture of the T1SS is closely related to the secretion system of the multidrug efflux pumps called resistance nodulation division (RND), which secretes most antibacterial molecules out of the cells, contributing to antibiotic resistance (Piddock, 2006). In contrast, the T2SS, including components called general secretion pathway (Gsp), ensures the transport of hydrolysing enzymes and toxins (Tauschek *et al.*, 2002; Kulkarni *et al.*, 2009; Nivaskumar and Francetic, 2014). Secretion of extracellular enzymes is often thought of as a ‘public goods’ service as they can provide an auxiliary function to bacteria otherwise lacking a given phenotype, for example the degradation of recalcitrant organic phosphorus or carbohydrate polymers (Lidbury *et al.*, 2017; Reintjes *et al.*, 2017; Enke *et al.*, 2019; Smith and Schuster, 2019).

In this study, our aim was to further investigate β-lactamase resistance in *E. coli* ST131 isolated from a UK river system downstream of a Waste Water Treatment Plant (WWTP), to establish the mechanism of enzyme secretion. (Wiles *et al.*, 2008; Gibreel *et al.*, 2012; Pitout, 2012; Nicolas-Chanoine *et al.*, 2014; Sarowska *et al.*, 2019; Whitmer *et al.*, 2019) We report that this strain provided a protective effect to a susceptible *E. coli* river isolate against cefotaxime. Further investigations demonstrated that CTX-M-15 was secreted and a role for T1SS was established.

## Results

### Phenotypic and genotypic testing of the two isolates

Resistance was determined for five antibiotics; strain 33 showed no phenotypic resistance to any of the antibiotics tested, in contrast, strain 48 was resistant to three of the five tested antibiotics (<u>Table S4</u>). Whole genome sequencing of strain 33 and 48 revealed the presence of three ESBLs genes of clinical relevance in strain 48 only; *bla*_TEM_, *bla*_OXA_ and *bla*_CTX-M-15_ with TEM and CTX-M the main types of ESBLs. PCR and sequencing confirmed the presence of the three ESBLs genes in strain 48.

### Strain 48 constitutively expresses a secreted β-lactamase

The two *E. coli* river isolates, 33 and 48, were challenged with cefotaxime (8 μg/ml, 16 μg/ml, 32 μg/ml and 64 μg/ml); growth of strain 33 was inhibited by all cefotaxime concentrations whilst strain 48 was completely resistant (Fig 1A). Previous genomic analyses revealed strain 48 is predicted to possess three ESBLs, encoded by *bla*_TEM_, *bla*_OXA_ and *bla*_CTX-M-15_. Biochemical analyses revealed that strain 48 possessed extracellular β-lactamase activity in either the presence or absence of cefotaxime (Fig 1B), although greatest activity was observed during growth on the highest concentration of this antibiotic. As expected, *E. coli* strain 33, which was sensitive to the β-lactams, showed no secreted β-lactamase activity (Fig 1B).

**Figure 1.**
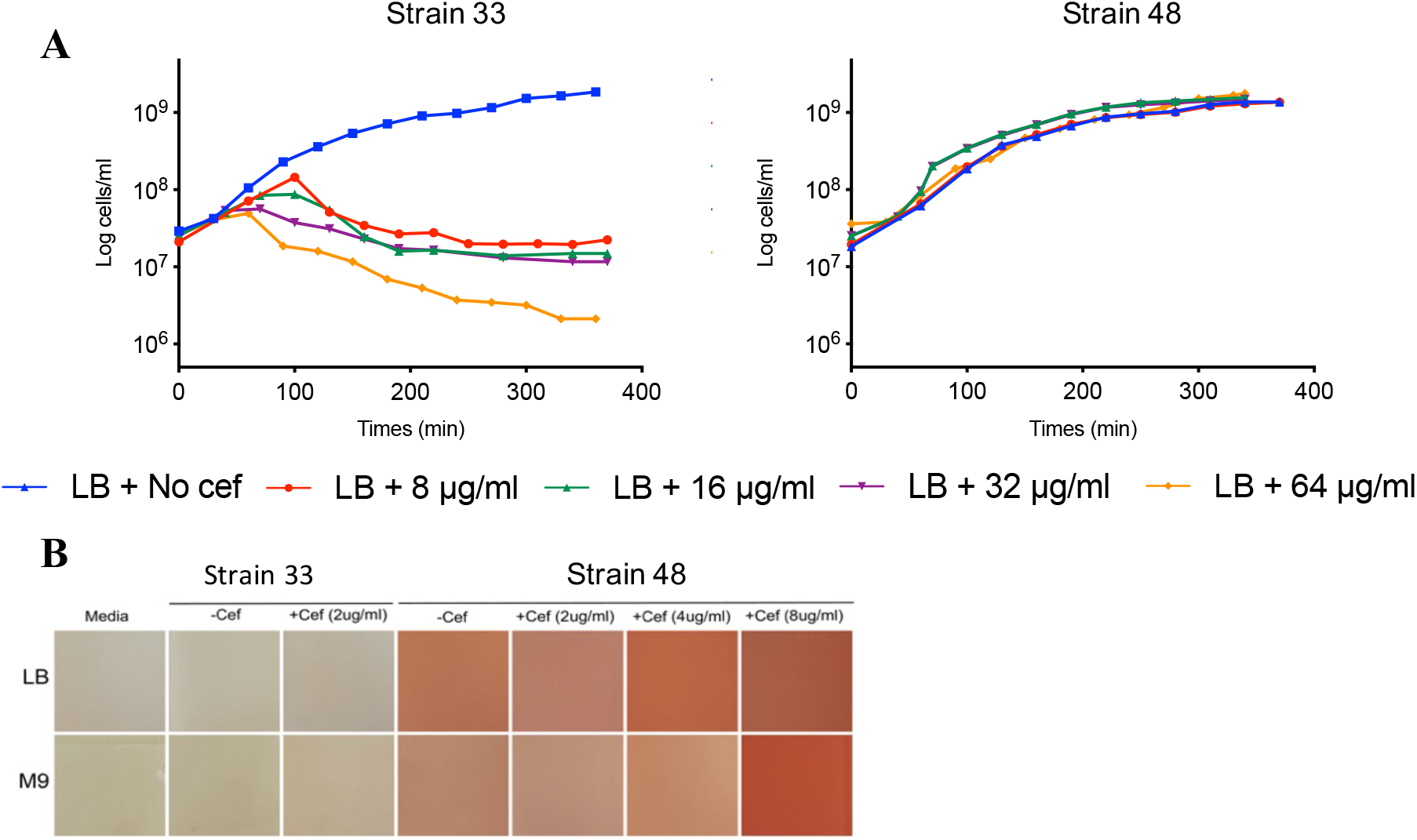
Secretion of β-lactamases by strain 48. (**A**) Growth of *E. coli* strains 33 and 48 cultivated in LB medium in varying cefotaxime concentrations, 8 μg/ml (in red), 16 μg/ml (in green), 32 μg/ml (in purple) and 64 μg/ml (in yellow). **(B)** Nitrocefin assay with various concentrations of cefotaxime (2 μg/ml, 4 μg/ml and 8 μg/ml) in presence of *E. coli* strain 33 or strain 48

### CTX-M-15 is responsible for conferring Cefotaxime resistance

To identify which of the three ESBLs was responsible for β-lactamase activity in strain 48, the proteome of this bacterium, partitioned into cellular (CP) and extracellular (XP) fractions, was analysed. Cells were grown in either the presence (8 μg/ml) or absence of cefotaxime. Both TEM and CTX-M-15 β-lactamases were identified in the CP and XP, however the relative abundance of these in the CP was ~10-fold lower than their relative abundance in the XP (Fig 2A). Exoproteomics identified 1845 proteins across all treatments, the majority of which represented a long tail of very low abundance proteins (<0.1%). Notably, CTX-M-15 was the third most abundant protein in the XP of strain 48, in either the presence (1.7%) or absence (2.48%) of cefotaxime (Fig 2A). The abundance of TEM in the XP was lower (absence, 0.38 %; presence 0.6%). Constitutive expression of CTX-M-15 was confirmed by RT-qPCR, which showed no significant difference in *bla*_CTX-M-15_ transcription in either the presence or absence of antibiotic (Figure S1).

**Figure 2.**
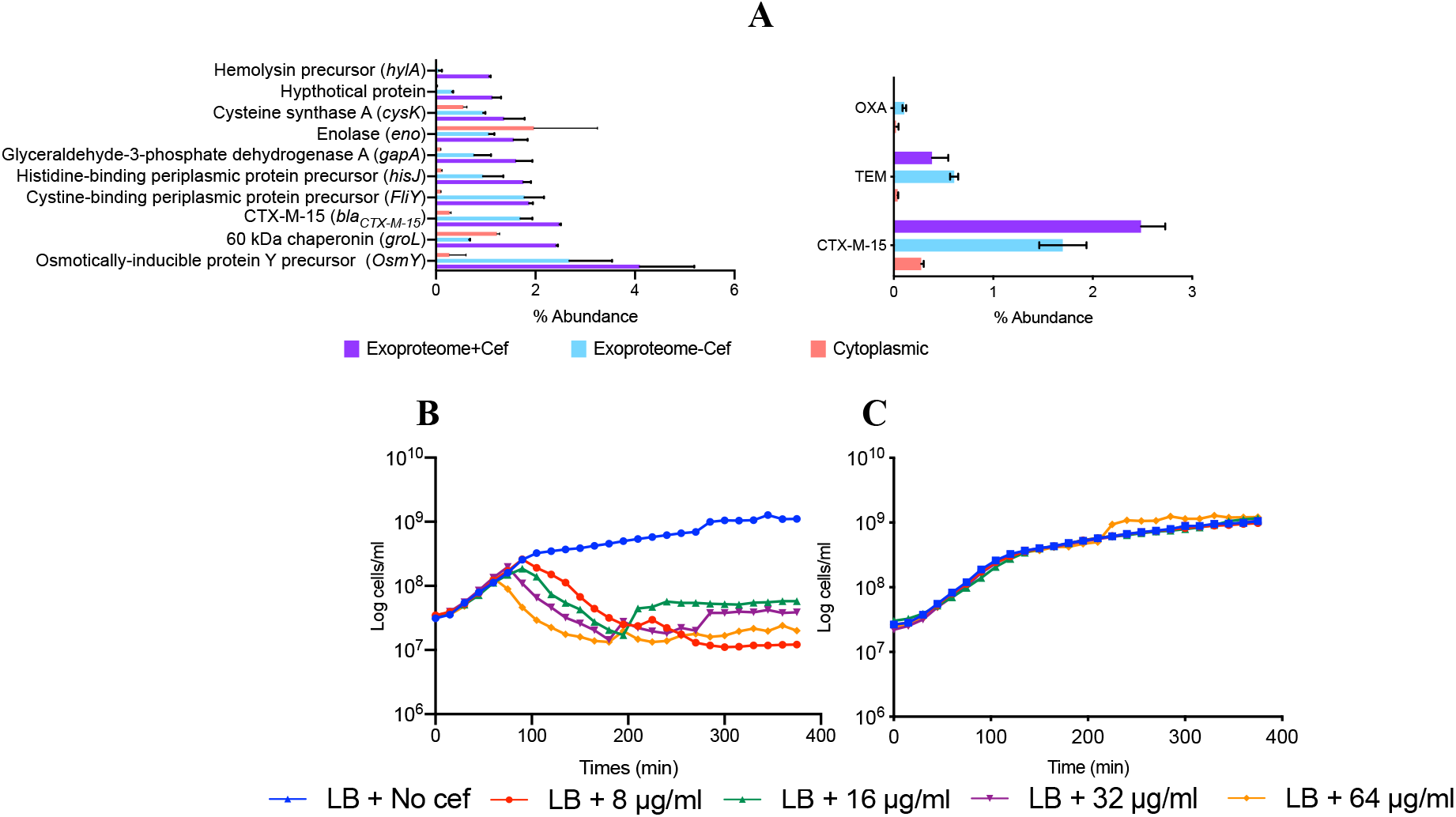
Identification of CTX-M-15 as the major secreted ESBL. (**A**) Top 10 of the most abundant proteins found in the exoproteome of strain 48 in presence of cefotaxime. Protein abundance was evaluated by MS/MS spectral counts which correlated linearly with the protein abundance. Error bars indicate mean of three replicates. Abundance of the three ESBLs proteins. The abundance of CTX-M-15 was significantly higher in presence of cefotaxime (p-value = 0.05). **(B)** Growth of JM109 and **(C)** JM109 CTX-M-15 cultivated in LB medium in varying cefotaxime concentrations 8 μg/ml (in red), 16 μg/ml (in green), 32 μg/ml (in purple) and 64 μg/ml (in yellow); JM109 empty plasmid.

To confirm if CTX-M-15 was responsible for conferring cefotaxime resistance, *bla*_CTX-M-15,_ *bla*_TEM_ and *bla*_OXA_ from strain 48 were separately cloned into the expression vector, pGEM-T. Plasmids were mobilised into a susceptible host, the commercial strain *E. coli* JM109, resulting in the strains JM109-OXA, JM109-TEM, and JM109-CTX-M-15. An empty vector control was also mobilised into JM109, creating the strain JM109-pGEM-T (Fig 2B). Only JM109-CTX-M-15 grew in the presence of cefotaxime confirming *bla*_CTX-M-15_ was essential for resistance to cefotaxime (Fig 2C).

### CTX-M-15 secretion provides protection to susceptible cells

Strain 33 was susceptible to cefotaxime, so to determine if secreted CTX-M-15 from strain 48 could complement a susceptible strain, strain 33 was grown in both presence and absence of cefotaxime in a conditioned medium (CM) used for growing strain 48. Strain 33 grew in CM in the presence of cefotaxime demonstrating that strain 48 secreted sufficient quantities of CTX-M-15 into the medium to degrade the antibiotic and prevent inhibition of the otherwise susceptible strain 33 (Fig 3A).

**Figure 3.**
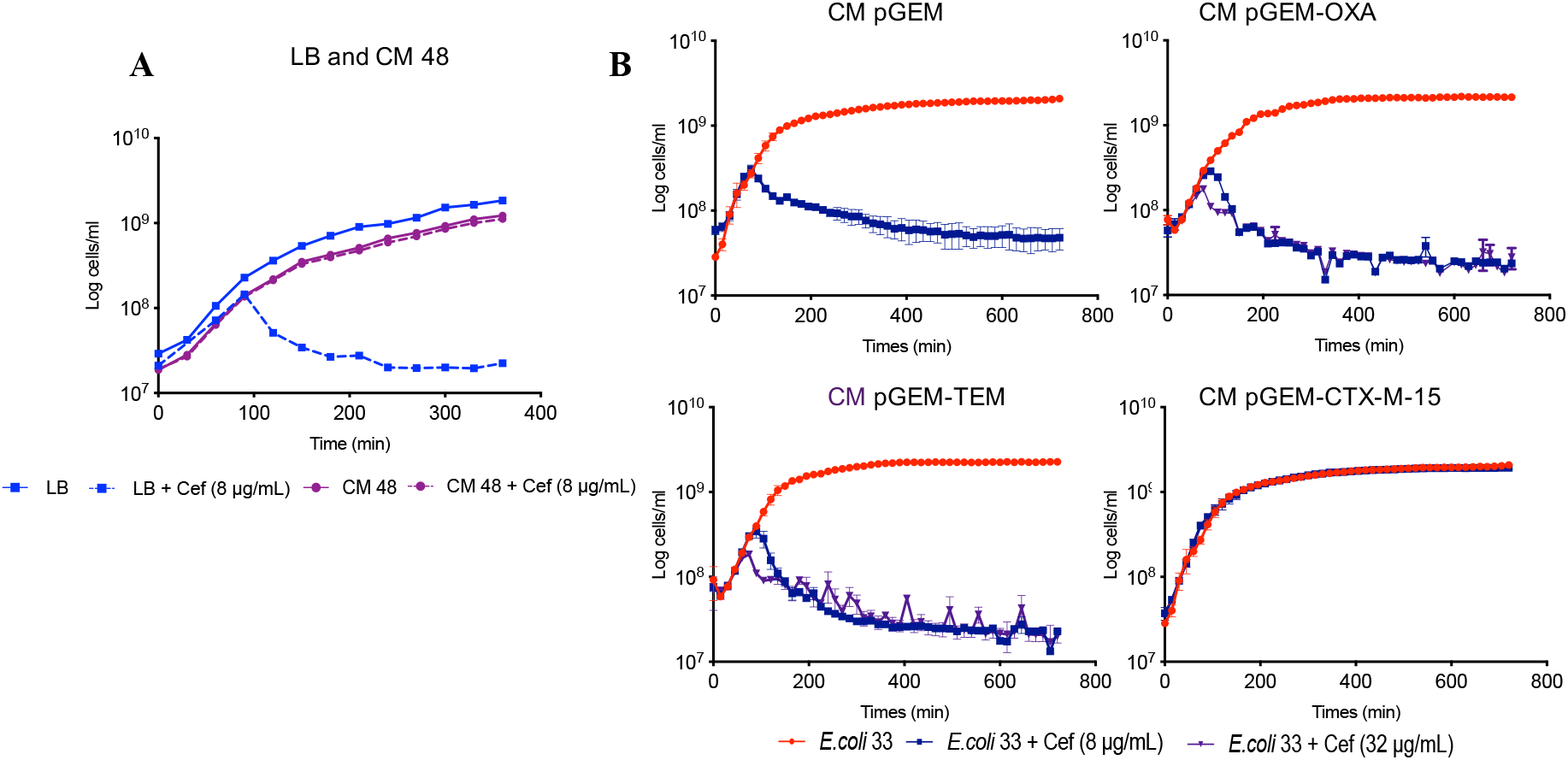
Protective effect of conditioned medium obtained from strain 48 growth chambers. (A) Growth of *E. coli* strain 33 (in blue) cultivated in fresh LB media compared to growth in CM of 48 (in purple). (B) Growth of strain 33 in CM of pGEM, CM of pGEM-OXA, CM of pGEM-TEM, CM of pGEM-CTX-M-15 in absence (in red) and presence of 8 μg/ml (in blue) and 32 μg/ml (in purple) of cefotaxime.

The protection given by the secreted CTX-M-15 of strain 48 was confirmed with the engineered JM109 strains. Only CM from JM109-CTX-M-15 facilitated the growth of strain 33, whilst CM from TEM and OXA producing strains did not (Fig 3B). Finally, we tested the stability of secreted CTX-M-15 by storing CM from JM109-CTX-M-15 at 4°C for 24 h, 48 h and 72 h, prior to inoculation with strain 33. Again, strain 33 grew in the presence of cefotaxime (<u>Fig S2</u>). Together, this demonstrated that CTX-M-15 is functionally stable after secretion outside of the cell and can provide protection to otherwise susceptible bacterial strains.

### The T1SS is involved in the secretion of CTX-M-15

To identify a mechanism of secretion for this CTX-M-15, we first investigated the role of OMVs, which have previously been reported to express ESBL activity (Kim *et al.*, 2018). To remove OMVs from the supernatant, CM obtained from strain 48 was additionally filtered through a 0.02 μm filter. However, the susceptible strain 33 still grew in the presence of cefotaxime when grown in conditioned medium. This suggests that OMVs either lysed during filtration or that they are not the main mechanism for CTX-M-15 secretion (Fig S3).

In agreement with their abundance in the XP of strain 48, *in silico* prediction revealed both CTX-M-15 and TEM contained the SP1 leader sequence required for translocation across the cytoplasmic membrane by the Sec pathway (Fig S4, Table S5). Strain 48 was predicted to contain four known secretion systems T1SS, T2SS, T4SS and T5SS, that were therefore candidates for CTX-M-15 secretion. T1SS and T2SS have potential to be involved in the secretion of hydrolytic enzymes, such as ESBLs. Mutant *E. coli* strains defective for genes required for either T1 secretion (*tolC*) or T2 secretion (*gspD*) were obtained from the Keio database, as was the parental wild type, *E. coli* BW 25113. pGEM-T-CTX-M-15, conferring cefotaxime resistance, was mobilised into all three strains (*E. coli* BW+, *E. coli ΔtolC*+, *E. coli ΔgspD*+) which were again subjected to growth in the presence of cefotaxime. In addition, all three strains, *E. coli* BW, *E. coli ΔtolC* and *E. coli ΔgspD* were transformed with an empty pGEM-T vector as controls. As expected, these three control strains failed to grow in presence of cefotaxime (Fig S5A). *E. coli* BW+, *ΔtolC*+ and *ΔgspD*+ all grew on varying concentrations of cefotaxime, but mutation of either T1SS or T2SS significantly inhibited their growth rates (Fig 4A, Fig S5B). For the *ΔgspD* mutant, one cefotaxime concentration (8 μg/ml) showed a significant (P<0.01) reduction in growth rate, whilst *ΔtolC+* displayed a significantly (P<0.05) slower growth rate when challenged with all three concentrations of cefotaxime (Fig 4A). To determine if this partial growth inhibition in either mutant was due to an inhibition by CTX-M-15 extracellular secretion, the growth of strain 33 on CM obtained from *E. coli* BW+, *E. coli ΔtolC*+ and *ΔgspD*+ cultures was monitored, in the presence or absence of cefotaxime. Similar to the growth of both secretion mutants, the growth rate of strain 33 was significantly reduced by the presence of cefotaxime when cultured in CM obtained from either *ΔtolC+* or *ΔgspD+* relative to the BW wild type CM (Fig 4B, Fig S5C). Mutation of *ΔtolC* (T1SS) resulted in a greater sensitivity to cefotaxime for either the producer or the susceptible strain, indicating this secretion system is involved in, but not essential for CTX-M-15 secretion. Given we also observed a smaller, but significant effect when mutating *gspD* (T2SS), it is likely that both these systems can facilitate CTX-M-15 secretion, with T1SS being the most important. Together, these data suggest secretion of CTX-M-15 is not a passive process and is facilitated by common secretion systems present in widespread bacteria.

**Figure 4.**
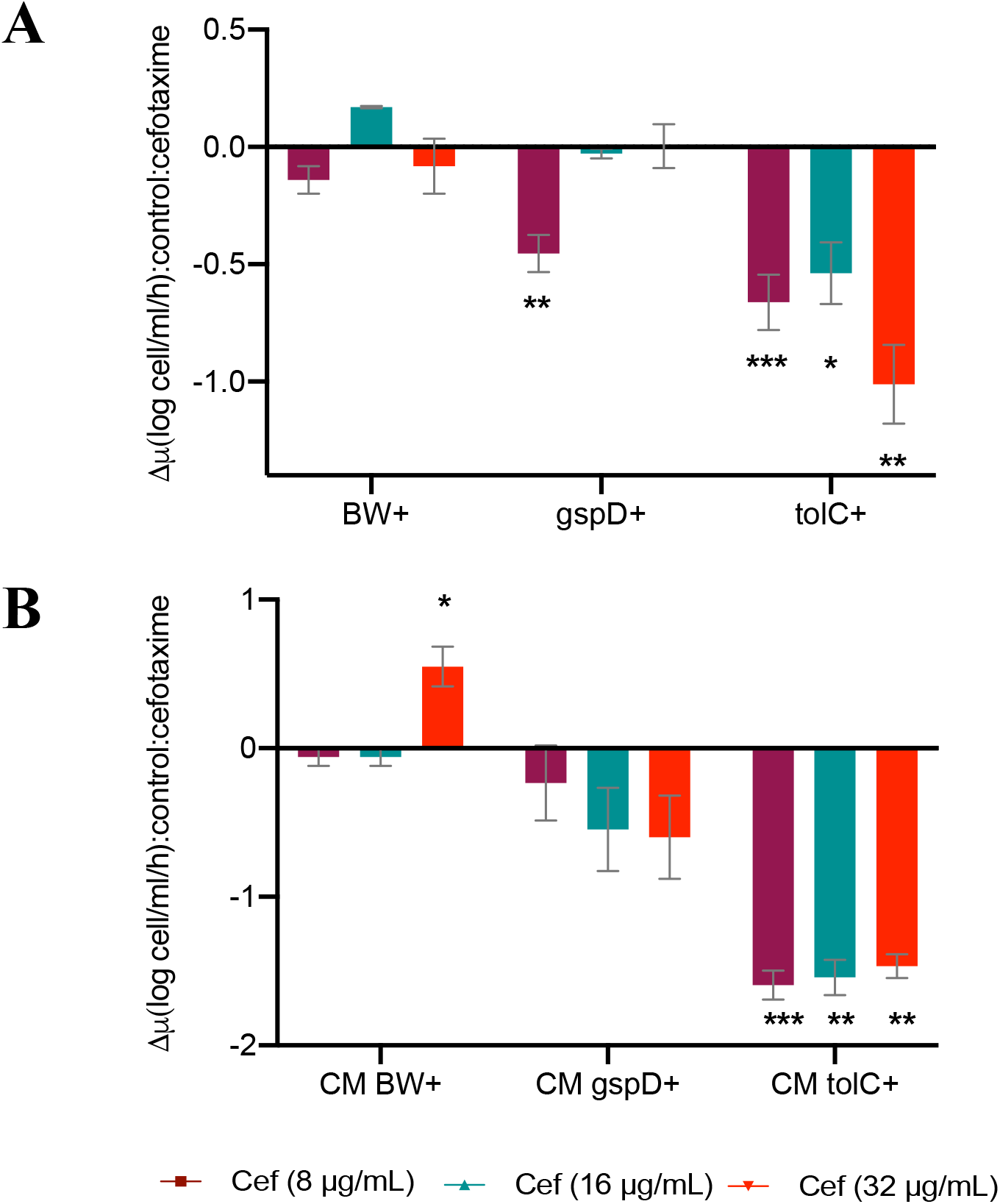
The potential role of T1SS in the secretion of CTX-M-15. (**A**) Growth rate (μ) of *E. coli* BW+, *E. coli* Δ*gspD+* and *E. coli* Δ*tolC+* in presence of 8 μg/ml (in magenta), 16 μg/ml (in green) and 32 μg/ml (in red) of cefotaxime. (**B**) Growth rate of strain 33 cultivated in CM obtained from *E. coli* BW+ (CM BW+), *E. coli* Δ*gspD*+ (CM Δ*gspD*+) and *E. coli* Δ*tolC*+ (CM Δ*tolC*+) in presence of 8 μg/ml (in magenta), 16 μg/ml (in green) and 32 μg/ml (in red) of cefotaxime. Graphs show the difference between the growth rate in the presence and the absence of antibiotic. The value indicates the mean ± standard deviation of three biological replicates. t-test, * significant at p-value < 0.05; ** significant at *p*-value < 0.01; *** significant at *p*-value < 0.001.

## Discussion

Secretion of ESBLs into the surrounding environment may reduce any damage to the cell wall by preventing entry of β-lactam into the periplasm where they would be degraded by enzymes such as OXA or TEM. It is also likely to be an important ecological trait, enabling otherwise susceptible bacteria to survive long enough to allow mobilisation of plasmid-encoded *bla* genes through conjugation (De La Cruz and Davies, 2000; Amos *et al.*, 2014). Whilst strain 48 possessed three annotated ESBLs, the data presented here proves that CTX-M-15 was the major secreted ESBL conferring resistance and providing a protective effect. This was further confirmed by exoproteomics, which revealed that CTX-M-15 was the third most abundant protein in the XP. The significant secretion of this resistance enzyme may well explain the rapid global dissemination of CTX-M-15 in clinical strains (Woerther *et al.*, 2013; Amos *et al.*, 2014; Bevan *et al.*, 2017; Amos *et al.*, 2018).

Whilst the presence of ESBLs in OMVs has been linked to extracellular degradation of β-lactam antibiotics (Ciofu *et al.*, 2000; Gonzalez *et al.*, 2016; Kim *et al.*, 2018) to the best of our knowledge, this is the first-time secretion of an individual ESBL (i.e CTX-M-15) has been linked to T1SS and proven to confer protection to susceptible bacteria. Our data clearly demonstrated CTX-M-15 was actively secreted into the extracellular milieu with no evidence that OMVs played a role. The ESBL NDM-1, which is the most widespread carbapenemase worldwide, has a lipobox proximal to its SP1 leader sequence that enables anchoring to the outer-membrane and subsequent export in OMVs (Gonzalez et al., 2016), a process known as lipidation. Removal of this lipobox inhibited NDM-1 export *via* OMVs and the enzyme accumulated in the periplasm. Lipidation only occurs in a few ESBLs, such as BRO-1 (from the human pathogen *Moraxella catarrhalis*) and PenA (from *Burkholderia pseudomallei*) (Bootsma *et al.*, 1996; Randall *et al.*, 2015), and does not include CTX-M-15, which may explain the lack of OMV involvement. Whilst OMVs were implicated in the secretion and extracellular functioning of a non-lipidated serine β-lactamase CTX-M-1 (Kim *et al.*, 2018), in our study removal of vesicles by membrane filtration (0.02 μm) from the culture supernatant did not reduce the efficacy of CTX-M-15 to confer a protective effect suggesting secretion occurred through an alternative mechanism. Even if some vesicles were disrupted during membrane filtration, our data proved that CTX-M-15 was actively secreted not associated with membranes.

In contrast, we discovered that mutation of key genes required for either Type-I or Type-II secretion inhibited the efficacy of CTX-M-15-induced protection, both to the producer and the susceptible strains. Interestingly, our data proved that T1SS played a role in the direct secretion of CTX-M-15, despite the fact that this ESBL contains a leader sequence for localisation in the periplasm. The T2SS may also play a role in secretion of CTX-M-15, although such a small amount of proteins in the XP could be explained by spontaneous release of the periplasmic proteins. Indeed, out of the top five most abundant proteins in the XP, OsmY and two ligand binding proteins were also predicted to be periplasmic (Yim and Villarejo, 1992; Oh *et al.*, 1994), yet were still found in the culture supernatant. Exoproteomics often captures a range of periplasmic proteins, especially ligand binding proteins (Christie-Oleza and Armengaud, 2010; Christie-Oleza and Armengaud, 2015; Lidbury *et al.*, 2016) which suggests that the outer membrane may be leaky. The mechanism of secretion for OsmY has not been studied but it was used in biotechnology to deliver proteins into the medium *via* C-terminal fusion (Qian *et al.*, 2008; Bokinsky *et al.*, 2011). Reports concerning secretion in *E. coli* remain elusive mainly because non-pathogenic laboratory strains generally express a small amount of proteins in the culture medium (Papanikou *et al.*, 2007; Kotzsch *et al.*, 2010). Identifying the causal mechanism for CTX-M-15 secretion could help develop novel therapeutic drugs to block secretion as we have proven it is essential for resistance under exposure to cefotaxime.

Despite not being able to clearly identify the secretory mechanism of strain 48, we showed that CTX-M-15 is the major secreted ESBL conferring a protective effect to neighbouring susceptible cells. It is feasible that secretion of CTX-M-15 represents an evolutionary advantage, as no damage would occur to the cell-wall if the antibiotic is disabled outside of the cell in opposition to hydrolysis in the periplasm. The role of environmental contamination in the transmission of *Enterobacteriaceae* and in particular *E. coli* ST131 is increasingly recognize. However, factors influencing duration of survival in the environment have not yet been extensively studied.

## Materials and methods

### Bacterial strains and growth medium

Bacterial strains used in this study are listed in the supplementary information (<u>Table S1</u>). Environmental *E. coli* strains were isolated from the River Sowe, Coventry, UK; namely *E. coli* ST3576 O8:H7 (strain 33) and *E. coli* ST131 O25:H4 (strain 48). Commercial laboratory strains of *E. coli* JM109, *E. coli* BW2511, *E. coli* JW5503 and *E. coli* JW5707 were also used. Cells were routinely grown in Luria Bertani (LB) liquid broth (10 g/L tryptone, 5 g/L yeast extract, 10 g/L sodium chloride) or LB agar medium (addition of 15 g/L agar). The following antibiotics were supplemented when required: 8 μg/ml of cefotaxime, 100 μg/ml of ampicillin, 5 μg/ml of kanamycin. Additionally, culture medium for the JM109 was supplemented with isopropyl β-D-thigalactosidase (IPTG) (0.1 M) and X-galactosidase (20 mg/ml) to induce expression of recombinant CTX-M-15. Cells were incubated at 37°C with either shaking (200 rpm) or static conditions.

### Antimicrobial phenotypic screening

Oxid™ antibiotic discs were used to determine phenotypic resistance profiles. Strain 33 and 48 were streaked on LB agar plates and discs containing either 25 μg ampicillin, 5 μg cefotaxime, 10 μg imipenem, 30 μg chloramphenicol, or 8 μg erythromycin were added on top of the plates. All the plates were incubated overnight at 37°C.

### ESBLs genotypic screening

Single colonies of strains 33 and 48 were picked and individually inoculated in to 10 ml LB and incubated overnight at 37°C with shaking at 150 rpm. Cultures were then centrifuged at (1500 rpm for 10 min) and supernatant discarded. Pellets were resuspended in 500 μl PBS and used for DNA extraction using the MPBio FastDNA™ spin kit following the manufacturer’s guidelines. Specific primers for amplification of ESBLs genes, *bla*_CTX-M-15_, *bla*_TEM_ and *bla*_OXA_ were designed from the Illumina sequencing done previously (Hill, 2016) (<u>Table S2</u>). PCR reactions were done using 12.5 μL Master Mix 2X (Promega), 1.25 μL DMSO, 0.8 μM forward primer, 0.8 μM reverse primer, 2 μL DNA template and dH_2_O for a final PCR reaction volume of 25 μl. PCR was performed at an initial denaturation temperature at 95°C for 5 min, followed by 34 cycles of denaturation at 95°C for 30 sec, annealing temperature (Ta) at 55-66°C (depending on the primer set) for 30 sec and extension for 1 min 50 sec. A final extension was performed at 72°C for 5 min.

### Antibiotic resistance screening

Growth curves were implemented in 96-well plates with 200 μl culture per well and incubated at 37°C with shaking (200 rpm) in a microplate reader (POLARstar Omega, BMG labtech). As inoculum, overnight starter cultures of each bacterial strain (5 ml) were diluted to an initial concentration of 3 × 10^7^ cells/ml. Culture media were supplemented with 0, 8, 16, 32 or 64 μg/ml of cefotaxime (VWR International Ltd). Cell proliferation was determined by measuring the optical density at 600 nm for 8 or 12 h every 15 min. Each condition was set up in triplicate. Exponential growth rates were calculated for the growth of *E. coli* BW+, *E. coli* ΔtolC+ and *E. coli* ΔgspD+ and for the protective effect on strain 33 by using the following formula; P(t) = P_0_e^rt^ where P(t) is the amount of cell number at time t, P_0_ the initial cell number, r the growth rate and t the number of periods (Hall et al., 2014). Two-sample t-Test was performed to compare the significance of the growth rate differences.

### Generation of conditioned medium

Strain 48, or the engineered laboratory *E. coli* strains harbouring CTX-M-15 were grown in the presence of cefotaxime (8 μg/ml). After overnight growth, cells were removed by pelleting (3228 × G for 15 min). Supernatant was carefully filtered through a 0.22 μm membrane (Fisher Scientific) to avoid cell lysis. Conditioned medium (CM) was diluted 1:1 (v/v) parts with fresh LB medium and supplemented with varying concentrations of cefotaxime. Overnight cultures of susceptible strain 33 were inoculated (1% v/v) in the conditioned medium and grown as described above.

### Cloning of the ESBLs, *bla*_CTX-M-15_, *bla*_TEM_ and *bla*_OXA_

A full list of primers used in this study are presented in Table S2 and the genes *bla*_CTX-M-15_, *bla*_TEM_ and *bla*_OXA_ were cloned from strain 48 into the cloning vector pGEM-T easy (Promega, UK) using the HiFi assembly kit (New England, Biolabs). The newly constructed plasmids pGEM-CTX-M-15, pGEM-TEM and pGEM-OXA, and a control empty-vector pGEM-T were transformed into *E. coli* JM109.

### Detection of β-lactamase activity by the nitrocefin assay

β-lactamase activity was assessed by colorimetric assay using the chromogenic cephalosporin compound nitrocefin (Thermo Scientific) (O’Callaghan et al., 1972). Strain 33 and strain 48 were inoculated in LB or M9 minimal media (33.9 g/L Na_2_HPO_4_, 15g/L KH_2_PO_4_, 5 g/L NH_4_Cl, 2.5 g/L NaCl, 20 % glucose, 1 M MgSO_4_, 1 M CaCl_2_) supplemented with 0, 2, 4 or 8 μg/ml cefotaxime. Strains were grown at 37°C until mid-exponential phase and supernatant was collected by first removing cells (4000 rpm for 15 min) and then gentle filtration through a 0.22 μm membrane (Fisher Scientific) to prevent cell lysis and removed intact cells. Supernatants were incubated with 15 μg/ml nitrocefin (stock concentration 500 μg/ml) at room temperature (~22°C) for 30 min.

### Determination of *bla*_CTX-M-15_ transcription in strain 48

Strain 48 was grown at 37°C in LB supplemented with 0 or 8 μg/ml cefotaxime. Diluted cultures were grown at 37°C with shaking (200 rpm) to exponential phase before RNA extraction. A detailed protocol for extraction, reverse transcription and quantitative PCR can be found in the supplementary methods.

### Preparation of exoproteome, total proteome samples and LC-MS/MS analysis

Exoproteomes and total proteomes of strain 48 were prepared by adapting the protocol described in Christie-Oleza and Armengaud (Christie-Oleza and Armengaud, 2010). Briefly, Strataclean beads (Agilent) were used to isolate proteins instead of TCA precipitation. A detailed procedure is provided in the supplementary material.

### Peptide identification and comparative proteomic analysis

A custom database was made with the genome of strain 48 by using Prokka v1.14.5 for annotation (Seemann, 2014) and MASCOT was used to assign peptide to protein by using the custom database, identified proteins were further analysed using Scaffold (Searle, 2010) (Protein threshold 99.9 %, minimum peptide 2, peptide threshold 80 %). The normalized spectral abundance factor (NSAF) (Zhu *et al.*, 2010) was calculated for each protein to compare the abundance for all proteins. Two-sample t-Test was used to determine if presence of antibiotic significantly impacted the proteins abundance.

### *In silico* prediction of protein localisation and secretion pathways

Analysis was done on the SignalP 5.0 server (http://www.cbs.dtu.dk/services/SignalP-5.0/) and enabled the prediction of the presence and the location of cleavage sites in the three β-lactamase proteins CTX-M-15, TEM, and OXA using the Fasta sequences generated in house (See supplementary information) (Almagro Armenteros et al., 2019). The TXSScan webtool (https://galaxy.pasteur.fr/) (Afgan et al., 2018) was used for prediction of the presence of secretion systems in strain 48 using the genome of *E. coli* 48.

## Acknowledgements

We gratefully acknowledge financial support from the Natural Environment Research Council (grant NE/N019857/1). SR was supported by a CENTA NERC studentship.

