## Supplemental Material for "Mechanisms involved in the active secretion of CTX-M-15 β-lactamase by pathogenic *E. coli* ST131"

### Materials and Methods

#### Bacterial strains and growth medium

**Table S1** Bacterial strains and plasmids used in this study

| Plasmid | Description | Reference/Source |
|---|---|---|
| pGEM-T | Commercial cloning vector, containing Amp$^R$ and T7 promoter | Promega |
| pGEM-CTX-M-15 | Derivative of pGEM-T, bearing the *bla*$_{CTX-M-15}$ resistance gene driven by a T7 promoter | This study |
| pGEM-TEM | Derivative of pGEM-T, bearing the *bla*$_{TEM}$ resistance gene driven by a T7 promoter | This study |
| pGEM-OXA | Derivative of pGEM-T, bearing the *bla*$_{OXA}$ resistance gene driven by a T7 promoter | This study |
| **Strain** | **Genotype and comments** | **References/Source** |
| *E. coli* JM109 | *endA1, recA1, gyrA96, thi, hsdR17 (rk–, mk+), relA1, supE44, Δ(lac- proAB), [F′ traD36, proAB, laqIqZΔM15]* | Promega |
| *E. coli* JM109 pGEM | *E. coli* JM109 bearing the pGEM vector | This study |
| *E. coli* JM109 CTX-M-15 | *E. coli* JM109 bearing the pGEM-CTX-15 vector | This study |
| *E. coli* JM109 TEM | *E. coli* JM109 bearing the pGEM-TEM vector | This study |
| *E. coli* JM109 OXA | *E. coli* JM109 bearing the pGEM-OXA vector | This study |
| *E. coli* BW 25113 | K-12 derivative *rrnB3* Δ*lacZ4787 hsdR514* Δ*(araBAD)567* Δ*9rhaBAD)568 rph-1* | (Datsenko and Wanner, 2000) |
| *E. coli* JW5503 Δ*tolC* | F-, *Δ(araD-araB)567*, *ΔlacZ4787*(::rrnB-3), *λ*$^-$, *ΔtolC732::kan*, *rph-1*, *Δ(rhaD-rhaB)568*, *hsdR514,* Δ*tolC* | (Baba *et al.*, 2006) |
| *E. coli* BW 25113 JW5707 ΔgspD | F-, *Δ(araD-araB)567*, *ΔlacZ4787*(::rrnB-3), *λ*$^-$, *ΔgspD735::kan*, *rph-1*, *Δ(rhaD-rhaB)568*, *hsdR514,* ΔgspD | (Baba *et al.*, 2006) |
| *E. coli* BW+ | Derivative of BW 25113 carrying pGEM-T -*bla*$_{CTX-M-15}$ | This study |
| *E. coli* Δ*tolC*+ | Derivative of BW 25113 carrying pGEM-T-*bla*$_{CTX-M-15}$ | This study |
| *E. coli* JW5707 ΔgspD+ | Derivative of BW 25113 carrying pGEM-T-*bla*$_{CTX-M-15}$ | This study |

### ESBLs genotypic screening

**Table S2** Primers used for PCR screening

| Target | ID | Sequence | Ta ($^{o}$C) | Size (bp) |
|---|---|---|---|---|
| *bla*$_{CTX\text{-}M\ 15}$ | CTX-M_15F | 5'-ATGGT AAAAATCACT CGCCAGTTC-3' | 63 | 876 |
| | CTX-M_15R | 5'-TTACA AACCG TCGGT GACGAT-3 | | |
| *bla*$_{TEM}$ | TEM_F | 5'-ATGCG CCTGG TAAGC AGAGT-3' | 55 | 1124 |
| | TEM_R | 5'-TTACC AATGC TTAAT CAGTG-3' | | |
| *bla*$_{OXA}$ | OXA_F | 5'-CAGTT ACTGG CGAAT GCAT-3' | 66 | 1287 |
| | OXA_R | 5'-CGTCC CGACT TGATT GAAG-3' | | |

### Determination of bla$_{CTX\text{-}M\text{-}15}$ transcription

Total RNA from strain 48 was obtained using the RNeasy Mini Kit (Qiagen), according to the manufacturer's instructions. Total RNA concentration was estimated using a NanoDrop ND-1000 spectrophotometer and the 260/280 and 260/230 ratios were examined to check for protein and solvent contamination. Integrity of RNA samples was confirmed by agarose gel. Contaminant DNA was removed using the TURBO DNA-free kit (Ambion). Total RNA was then reverse transcribed using the SuperScript II Reverse Transcriptase (Invitrogen), following manufacturer's guidelines. A control omitting the reverse transcriptase was carried out for each RNA sample to rule out residual genomic DNA contamination. The obtained cDNA samples were then used as qPCR templates. Expression of *bla*$_{CTX\text{-}M\text{-}15}$ was assessed using the cycle threshold (Ct) values normalized relative to the housekeeping gene *hcaT*, encoding for a 3-phenylpropionic transporter (Zhou *et al.*, 2011). Primers used for *bla*$_{CTX\text{-}M\text{-}15}$ and *hcaT* can be found in Table S3. qPCR reactions mixes were prepared as follows: 12.5 µl SYBR® GREEN master mix (Invitrogen), 0.4 µM forward primer, 0.4 µM reverse primer, 0.5 mg/ml BSA, 8.25 µL $dH_2O$, and 1 µl DNA template. Reactions were carried out in the 7500 Fast Real-Time PCR System (Applied Biosystem), with the following conditions: (1) initial denaturation at 95$^{o}$C for 10 mins, (2) denaturation at 95$^{o}$C for 15 sec, (3) annealing at 60$^{o}$C for 1 min. Step 2 and 3 were repeated for 40 cycles.

**<u>Table S3</u>** Target probes used for quantification in the qPCR

| Target | Name | Sequence | Reference |
|---|---|---|---|
| *blaCTX-M-15* | CTX-M_15F | 5’-ATGGTAAAAATCACTCGCCAGTTC-3’ | This study |
| | CTX-M_15R | 5’-TTACA AACCG TCGGT GACGAT-3’ | |
| *hcaT* | hcaT_F | 5’-GCTGCTCGGCTTTCTCATCC-3’ | (Zhou *et al.*, 2011) |
| | hcaT_R | 5’-CCAACCACGCTGACCAACC-3’ | |

**Preparation of exoproteome, total proteome samples and LC-MS/MS analysis**

Exoproteomes and total proteomes of strain 48 were prepared as previously described in Christie-Oleza and Armengaud (2010) (Christie-Oleza and Armengaud, 2010), with modifications; cells (50 ml) were grown until mid-exponential phase in M9 minimal medium at 37°C with shaking (200 rpm) and then centrifuged (4,000 rpm for 15 min) at 4°C. Exoproteome: Supernatants were then filtered through two consecutive low-proteins-binding FisherBrand™ sterile PVDF filters (0.45 µm filter, followed by 0.22 µm filter) (Fisher Scientific) and acidified at pH 5 using a solution of trifluoroacetic acid 10% (v/v). 50 ml of supernatant was incubated overnight on a rotor wheel (40 rpm/min) with 30 µl of Strataclean beads (Agilent). Beads were collected by centrifugation (2,000 rpm for 1 min) and resuspended in lithium dodecyl sulfate (LDS) (Expedeon Ltd) amended with the reducing agent, dithiothreitol (DTT) (Expedeon Ltd). Sample were heated at 90°C for 5 min and then sonicated for 5 min. This cycle was repeated twice.

Total proteome: Cell pellets were resuspended in 3.5 ml of TrisHCl pH 7.8 20 mM and lysed using the French Pressure Cell Press (3 cycles at 1000 Pa, cooling samples in ice in between cycles).

LC-MS sample preparation: The protein pellets were resuspended in LDS with 1 % β-mercaptoethanol. Purified proteins were separated on a 1D-SDS PAGE (Expedeon Ltd) by protein migration performed at 180 V for 40 min. Following protein separation, the obtained gel was stained using SimplyBlue™ SafeStain (ThermoFisher). The stained protein gels were washed three times with $dH_2O$ and then left to distain overnight. Each gel band was then excised and transferred into a sterile microcentrifuge tube. In-gel reduction and alkylation of the proteins were performed using dithiothreitol and iodoacetamide, respectively. Proteins were digested overnight with 40 µl of trypsin solution (2.5 ng/µl) and peptides were recovered using a formic acid/acetonitrile extraction buffer prior to analysis using a nano LC-ESI-MS/MS Ultimate 3000 LC system (Dionex-LC Packings) associated to an Orbitrap Fusion mass spectrometer (Thermo Scientific).

### *In silico* prediction of protein localisation and secretion pathways

> CTX-M-15 [*E. coli* 48]

MVKKSLRQFTLMATATVTLLLGSVPLYAQTADVQQKLAELERQSGGRLGVALINTA
DNSQILYRADERFAMCSTSKVMAAAAVLKKSESEPNLLNQRVEIKKSDLVNYNPIAE
KHVNGTMSLAELSAAALQYSDNVAMNKLIAHVGGPASVTAFARQLGDETFRLDRT
EPTLNTAIPGDPRDTTSPRAMAQTLRNLTLGKALGDSQRAQLVTWMKGNTTGAASI
QAGLPASWVVGDKTGSGGYGTTNDIAVIWPKDRAPLILVTYFTQPQPKAESRRDVL
ASAAKIVTDGL

> TEM [*E. coli* 48]

MSIQHFRVALIPFFAAFCLPVFAHPETLVKVKDAEDQLGARVGYIELDLNSGKILESF
RPEERFPMMSTFKVLLCGAVLSRVDAGQEQLGRRIHYSQNDLVEYSPVTEKHLTDG
MTVRELCSAAITMSDNTAANLLLTTIGGPKELTAFLHNMGDHVTRLDRWEPELNEAI
PNDERDTTMPAAMATTLRKLLTGELLTLASRQQLIDWMEADKVAGPLLRSALPAG
WFIADKSGAGERGSRGIIAALGPDGKPSRIVVIYTTGSQATMDERNRQIAEIGASLIKH
W

> OXA [*E. coli* 48]

MLAVKIKPFTKPILIMKNTIHINFAIFLIIANIIYSSASASTDISTVASPLFEGTEGCFLLY
DASTNAEIAQFNKAKCATQMAPDSTFKIALSLMAFDAEIIDQKTIFKWDKTPKGMEI
WNSNHTPKTWMQFSVVWVSQEITQKIGLNKIKNYLKDFDYGNQDFSGDKERNNGL
TEAWLESSLKISPEEQIQFLRKIINHNLPVKNSAIENTIENMYLQDLDNSTKLYGKTGA
GFTANRTLQNGWFEGFIISKSGHKYVFVSALTGNLGSNLTSSIKAKKNAITILNTLNL

### Results

**Table S4** Phenotypic resistance profile of strain 33 and 48

| Isolate | Ampicillin (25 μg) | Cefotaxime (5 μg) | Imipenem (10 μg) | Chloramphenicol (30 μg) | Erythromycin (8 μg) |
|---|---|---|---|---|---|
| 33 | No | No | No | No | No |
| 48 | Yes | Yes | No | No | Yes |

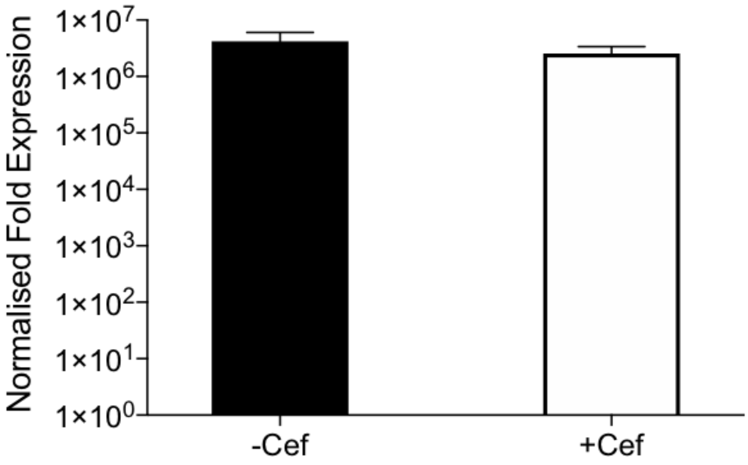


**Figure S1** Expression of *bla*$_{CTX\text{-}M\text{-}15}$ in strain 48 with and without cefotaxime. No significant difference in transcription following the addition of cefotaxime (8 μg/ml) (p-value=0.3).

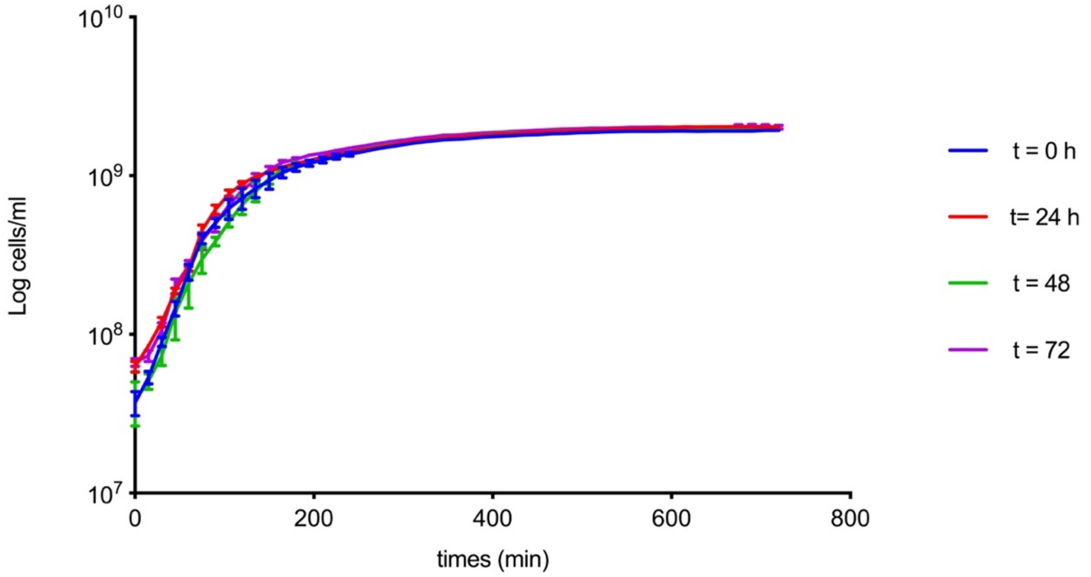


**Figure S2** Growth of strain 33 cultivated in CM of pGEM-CTX-M-15 in presence of 8 µg/ml of cefotaxime. Experiment was conducted with the CM pGEM-CTX-M-15 being at 4ºC for 24h (in blue), 48h (in red), and 72h (in green).

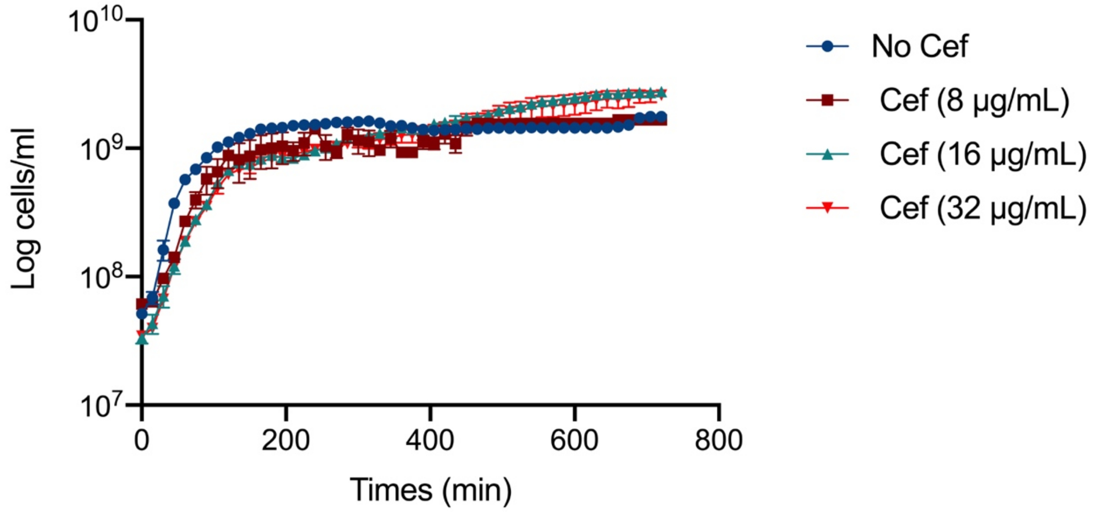


**Figure S3** Growth of *E. coli* strain 33 cultivated in CM obtained from *E. coli* 48 after filtration of the CM through a 0.02 µm. Experiment was conducted in absence (in blue) and presence of e 8µg/ml (in dark red), 16µg/ml (in green) and 32 µg/ml (in red) of cefotaxime.

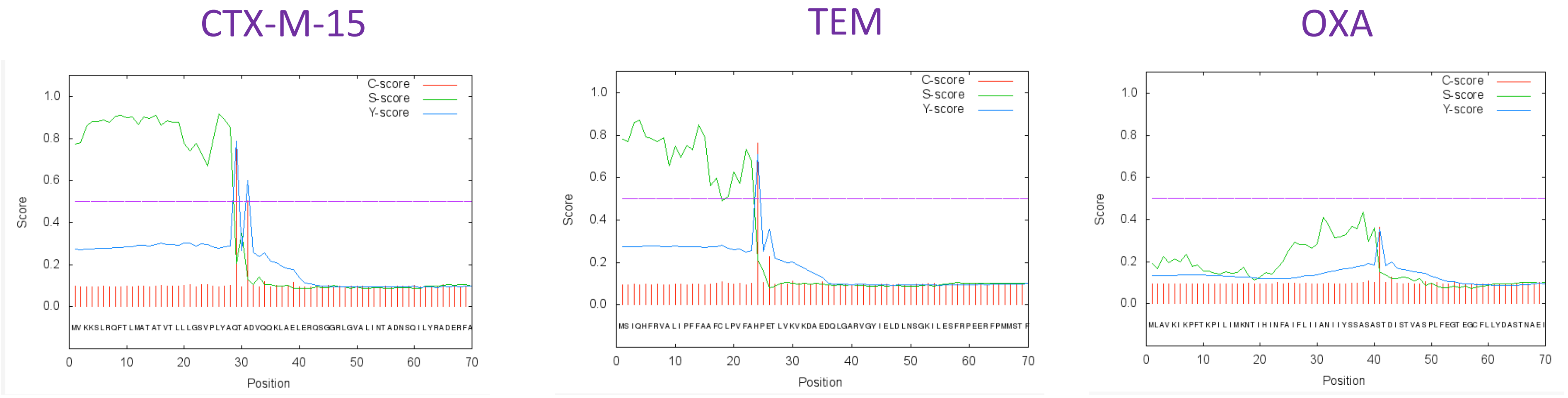


**Figure S4** SignalP server prediction for a signalling peptide in CTX-M-15 and TEM. The S score (green line) was used to predict the residues of the signal peptide. The cleavage site was determined by the C score (red line). The better cleavage site was determined by the Y score, which combines the predicting of the S and C scores.

**Table S5** D-Score predicted by SignalP server

| | D-score (cut-off =0.570) |
|---|---|
| CTX-M-15 | 0.817 |
| TEM | 0.710 |
| OXA | 0.308 |

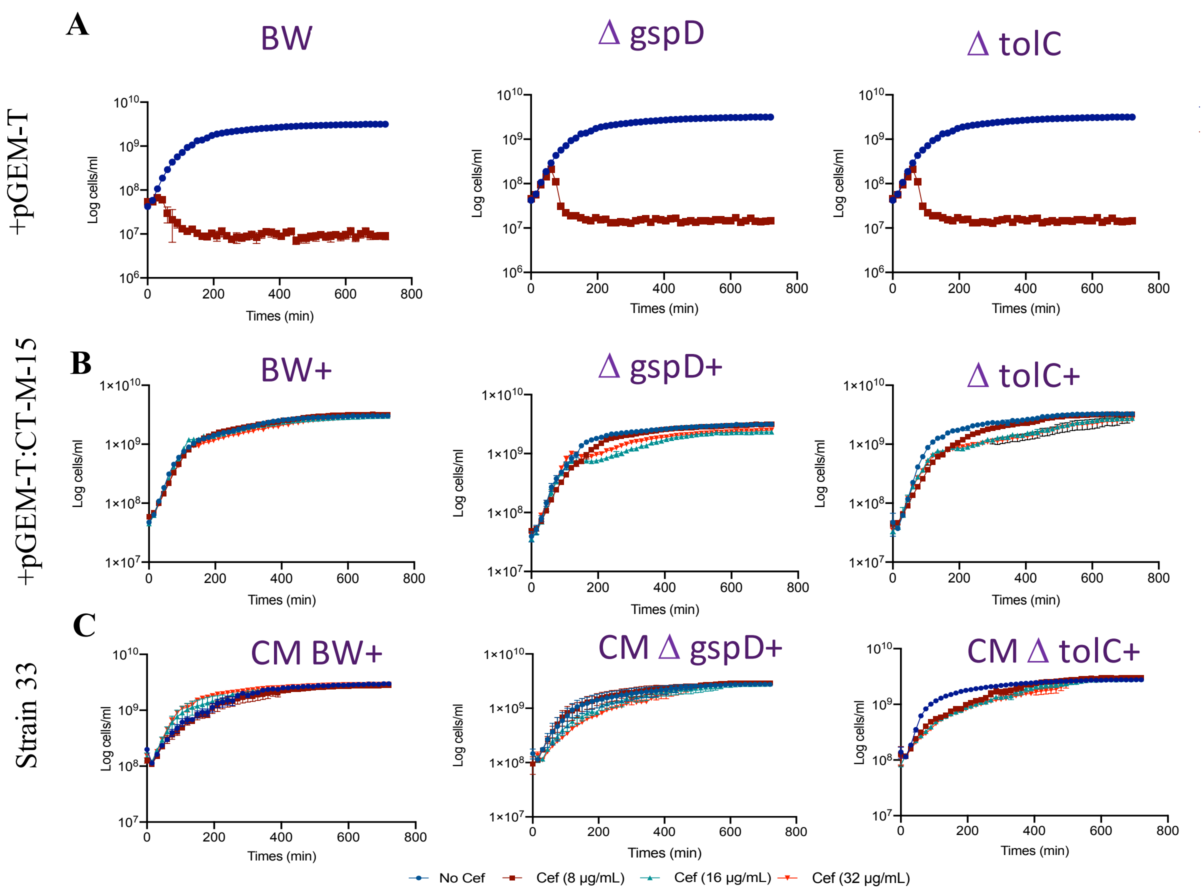


**Figure S5. The potential role of T1SS in the secretion of CTX-M-15. (A)** Growth of *E. coli* BW, *E. coli* Δ*gspD* and *E. coli* Δ*tolC* in absence (in blue) and presence of cefotaxime (8 µg/ml, in dark red). **(B)** Growth of *E. coli* BW+, *E. coli* Δ*gspD*+ and *E. coli* Δ*tolC*+ in absence (in blue) and presence of e 8µg/ml (in dark red), 16µg/ml (in green) and 32 µg/ml (in red) of cefotaxime. **(C)** Growth of strain 33 cultivated in different CM obtained from *E. coli* BW+ (CM BW+), *E. coli* Δ*gspD*+ (CM Δ*gspD*+) and *E. coli* Δ*tolC*+ (CM Δ*tolC*+) in absence (in blue) and presence of e 8µg/ml (in dark red), 16µg/ml (in green) and 32 µg/ml (in red) of cefotaxime

**Table S6** Exponential growth rate for *E. coli* BW+, *E. coli ΔtolC*+ and *E. coli ΔgspD*+. (t-Test; p=0.05).

| | Cefotaxime (µg/ml) | Exponential growth rate: replicate | | | *p*-value |
|---|---|---|---|---|---|
| | | 1 | 2 | 3 | |
| *E. coli* BW+ | 0 | 0.0289 | 0.0289 | 0.0242 | / |
| | 8 | 0.0254 | 0.0232 | 0.0232 | $2.28 \times 10^{-1}$ |
| | 16 | 0.0281 | 0.0288 | 0.0271 | $3.36 \times 10^{-1}$ |
| | 32 | 0.0245 | 0.0252 | 0.0252 | $4.30 \times 10^{-1}$ |
| *E. coli ΔgspD*+ | 0 | 0.0318 | 0.0274 | 0.0281 | / |
| | 8 | 0.0224 | 0.0217 | 0.0205 | **$1.43 \times 10^{-2}$** |
| | 16 | 0.0308 | 0.0273 | 0.0278 | $8.03 \times 10^{-1}$ |
| | 32 | 0.0308 | 0.03 | 0.0267 | $9.73 \times 10^{-1}$ |
| *E. coli ΔtolC*+ | 0 | 0.0316 | 0.0345 | 0.0324 | / |
| | 8 | 0.0231 | 0.0205 | 0.0218 | **$6.49 \times 10^{-4}$** |
| | 16 | 0.0252 | 0.0264 | 0.0200 | **$2.50 \times 10^{-2}$** |
| | 32 | 0.0193 | 0.0152 | 0.0134 | **$3.24 \times 10^{-3}$** |

*Significant p-values are denoted in bold.*

**Table S7** Exponential growth rate for strain 33 in presence of CM *ΔgspD*+ and CM *ΔtolC*+ (t-Test; p=0.05).

| | Cefotaxime (µg/ml) | Exponential growth rate: replicate | | | *p*-value |
|---|---|---|---|---|---|
| | | 1 | 2 | 3 | |
| *E. coli* BW+ | 0 | 0.0191 | 0.0175 | 0.0125 | |
| | 8 | 0.0166 | 0.0167 | 0.0128 | $7.01 \times 10^{-1}$ |
| | 16 | 0.0105 | 0.0222 | 0.0257 | $5.80 \times 10^{-1}$ |
| | 32 | 0.0288 | 0.0233 | 0.0245 | **$2.42 \times 10^{-2}$** |
| *E. coli ΔgspD*+ | 0 | 0.0134 | 0.0200 | 0.0290 | |
| | 8 | 0.0142 | 0.0094 | 0.0271 | $6.05 \times 10^{-1}$ |
| | 16 | 0.0104 | 0.0118 | 0.0129 | $1.85 \times 10^{-1}$ |
| | 32 | 0.0091 | 0.0105 | 0.0129 | $1.66 \times 10^{-1}$ |
| *E. coli ΔtolC*+ | 0 | 0.0377 | 0.0331 | 0.0351 | |
| | 8 | 0.0085 | 0.0076 | 0.0101 | **$4.08 \times 10^{-4}$** |
| | 16 | 0.0092 | 0.0102 | 0.0094 | **$2.81 \times 10^{-3}$** |
| | 32 | 0.0114 | 0.0106 | 0.0106 | **$3.08 \times 10^{-3}$** |

*Significant p-values are denoted in bold.*
